# Semantic predictability selectively modulates delta-band syntax tracking: evidence for good-enough parsing

**DOI:** 10.1101/2024.11.01.621536

**Authors:** M. Blake Rafferty, Edward C. Brown, Devin M. Casenhiser

## Abstract

Neural populations exhibit low-frequency phase synchrony to inferred phrase and constituent boundaries during speech comprehension, widely interpreted as reflecting the endogenous construction of syntactic structure. Good-enough processing (GEP) accounts suggest this structure-building is not obligatory for every utterance but graded: listeners may scale back syntactic computation when meaning can be recovered from local lexical-semantic context. Whether this graded reliance on syntax is reflected uniformly in delta-band neural activity remains unresolved, partly because syntactic processing comprises several dissociable demands that prior work has not modeled concurrently. We reanalyzed an open-source EEG dataset recorded during audiobook listening and used delta-band temporal response functions to model four syntactic constructs concurrently: syntactic surprisal, closing-node count, and integration and storage cost from Dependency Locality Theory. For each construct, we tested whether its contribution to the delta-band response was modulated by local semantic dissimilarity, computed from contextualized word embeddings. Semantic dissimilarity significantly modulated the response to syntactic surprisal and closing-node count, showed a smaller but reliable modulation of storage cost, and showed no modulation of integration cost; a repeated-measures comparison confirmed this graded hierarchy differed reliably across constructs. These results indicate that semantic predictability does not discount syntactic processing across the board but selectively affects the anticipation and construction of structure and its ongoing maintenance, while leaving retrieval of an already-resolved dependency largely unaffected. We interpret this pattern as sharpening, rather than merely reiterating, the claim that listeners prioritize efficiency during real-time comprehension.

## Introduction

Speech comprehension requires the human brain to encode linguistic representations from continuous auditory input under real-time constraints. Lower-level units such as phonemes and words may be identified via physiological correlates in the speech signal (Doelling et al. 2014), but comprehension beyond single words requires a listener to compute inter-word relationships, licensing their combination into multi-word syntactic units. Because reliable cues to these relationships are not always present in the signal itself, listeners must often infer them from previously encountered patterns and construct them endogenously (Meyer et al., 2020; Martin et al., 2020).

A substantial body of psycholinguistic work suggests this endogenous computation is not compulsory in the way it is sometimes assumed to be. Good-enough processing (GEP) accounts argue that listeners do not always construct a full, detailed syntactic parse of an utterance, but instead scale back the depth of syntactic analysis when meaning can be recovered through lexical-semantic or world knowledge. Doing so allows them to prioritize efficient interpretation over syntactic completeness (Ferreira et al., 2002; Ferreira & Patson, 2007; Karimi & Ferreira, 2016). Behavioral support for this view is well established by the examination of comprehension errors and reading-time studies in which readers accept implausible or ungrammatical structures when semantic plausibility offers an easy interpretive shortcut (Christianson et al., 2001; Sanford & Sturt, 2002). What remains underexplored in the research is whether, and how, this graded reliance on syntax is reflected directly in the neural computations that support structure-building.

We suggest that examination of low-frequency neural synchrony in the delta band (< 3 Hz) may well shed light on the neural computations involved in structure building vis a vis good enough processing. A wealth of electrophysiological evidence indicates that delta-band activity tracks the endogenous construction of syntactic structure during comprehension (Ding et al., 2016, 2017; Meyer et al., 2017; Kaufeld et al. 2020; Burroughs et al., 2021; Coopmans et al., 2022; 2025; ten Oever et al., 2023; Rafferty et al., 2023; 2024; for a review see Molinaro & Lizarazu, 2024), aligning with inferred phrase and constituent boundaries in a way that is dissociable from the prosodic and rhythmic cues with which those boundaries often coincide (Coopmans et al., 2025; Rafferty et al., 2023; Rafferty et al., 2026; cf. Glushko et al., 2022). Critically, this tracking appears to be genuinely syntactic rather than a general marker of composition or predictability: it depends on the presence of hierarchical phrase structure rather than on rhythmically recurring semantic information alone (Lu et al., 2022). It even persists when familiar words are replaced with pseudowords, provided listeners retain the cues needed to infer phrase and thematic structure (Rafferty et al., 2024; cf., Coopmans et al., 2022).

At the same time, this alignment is not fixed: the features of structure that delta activity tracks appear to depend on the lexical, statistical, and structural context of the input, and potentially on language- or context-specific processing strategies. Recent work shows that delta-band tracking is better explained by top-down rather than by integratory parsing strategies in Dutch, which is a head-final language (Coopmans et al., 2025). Moreover, delta oscillations can be reshaped by the presence of sentence-level context even for lexical features such as word frequency (Slaats et al., 2024), and joint models that contain both structural and statistical predictors explain more of the delta-band response than the sum of what each explains on its own (Weissbart & Martin, 2024; Slaats & Martin, 2025). Delta-band tracking thus provides a signature that is at once genuinely syntactic and demonstrably sensitive to the lexical and statistical context in which structure is built, making it well suited to ask whether it is also shaped by semantic predictability in the way GEP would predict.

Despite this wealth of evidence that delta-band syntactic tracking is modulated by numerous properties of the input, the specific question raised by GEP, namely whether local semantic predictability directly modulates the neural cost of syntactic processing, has not yet been directly characterized. The closest precedent comes from work asking whether distributional, word-level probability interacts with syntactic structure in delta-band dynamics, motivated by a statistical-learning account in which distributional regularities serve as a cue to structure-building (Slaats et al., 2024). This work, along with a broader debate over whether hierarchical structure or sequential probability better explains processing demands during comprehension (Frank & Bod, 2011; Frank & Christiansen, 2018; Brennan & Hale, 2019), converges on the conclusion that syntactic and distributional information are separately encoded and jointly, rather than competitively, shape the neural signal (Roark et al., 2009; Nelson, et al., 2017). Indeed, syntactic and statistical predictors have been shown to interact synergistically in shaping neural dynamics more broadly (Weissbart & Martin, 2024), and delta-band responses to syntactic constituent closure have been shown to shift in timing depending on lexical surprisal (Slaats et al., 2024).

The present study extends this logic in two respects. First, rather than asking whether lexical, distributional probability interacts with syntax, we ask whether the predictability of the *semantics* of a word given its local context—rather than the frequency with which the word (i.e., the token) occurs in the context—interacts with syntax. Critically, this question is motivated not by a statistical-learning account of cue validity but by GEP’s specific claim that listeners scale back structural computation when semantic context can support interpretation. Second, rather than testing this interaction against a single composite syntactic index, we test it against four dissociable syntactic constructs concurrently, allowing us to not only ask whether such an interaction exists but also to determine in which components of syntactic processing it is concentrated.

Syntactic processing is not monolithic: computational and psycholinguistic accounts decompose it into dissociable demands, including the cost of predicting upcoming structure before it arrives, the cost of provisionally maintaining unresolved dependencies in memory, the cost of retrieving and integrating a dependent once its head becomes available, and the local structural work of completing a constituent (Gibson, 1998; 2000; Hale, 2001). A neural extension of GEP that treats these demands as a single construct cannot distinguish a general down-weighting of syntactic effort from a more selective one, in which only some components of syntactic processing, such as prediction and structure-building, are discounted under high semantic predictability while others, such as memory retrieval, are not.

In the present study, we addressed this question directly by modeling four syntactic constructs concurrently within a single delta-band temporal response function (TRF) framework: syntactic surprisal, derived from a probabilistic constituency grammar to index prediction over hierarchical structure specifically; closing-node count, indexing the local structural work of completing a constituent; and integration and storage cost, derived from dependency locality theory to index retrieval and maintenance demands during dependency formation respectively. We chose a structurally-derived measure of surprisal rather than one based on a lexical language model because surprisal computed over word sequences is sensitive to any latent factor that shapes word choice, including frequency and pragmatic context alongside structure, making it difficult to attribute a modulation of such a measure specifically to syntax (Slaats & Martin, 2025; Slaats et al., 2023). Surprisal derived from a probabilistic constituency grammar is comparatively less entangled with these factors: it is computed from the probabilities of the grammatical rules used to build a word’s syntactic structure rather than from surface word-sequence statistics. Consequently, it weights structural configuration more heavily than lexical identity or frequency. We do not claim this measure isolates syntax from lexical statistics entirely, since the two are not fully separable in natural language, but relative to lexical surprisal it shifts the balance toward structural predictability, making a modulation of the measure more readily interpretable in terms of syntactic rather than purely lexical processing.

Semantic predictability was operationalized as contextual dissimilarity, a measure introduced by Broderick and colleagues (2018) to index the semantic relationship between a word and its preceding context. For each content word, dissimilarity is computed as one minus the similarity between that word’s semantic representation and the average representation of the preceding words in its sentence, such that a word semantically congruent with its context yields low dissimilarity and a word that departs from it yields high dissimilarity. Broderick and colleagues showed that this measure elicits a centro-parietal negativity resembling the N400 that is diminished for time-reversed, unattended, and unintelligible speech, indicating that it tracks the semantic processing of comprehended speech rather than a purely distributional property of the stimulus. Subsequent work further showed that it is only weakly correlated with lexical surprisal and captures a dissociable, semantic-level contribution to the neural response (Broderick et al., 2018; 2019; 2022). In the present study, we used this measure as a continuous index of how predictable each word was given its semantic context and tested whether that predictability modulated the neural response to each of the four syntactic constructs.

These considerations yield two contrasting predictions for how semantic predictability should relate to the neural cost of syntactic processing. If GEP reflects a general, undifferentiated strategy of down-weighting syntactic effort whenever local semantics can carry the interpretive burden, then semantic predictability should modulate all four syntactic constructs comparably: prediction, structural completion, memory maintenance, and retrieval should each show a reduced neural response as semantic dissimilarity decreases. This would suggest that the brain treats syntactic computation as a collection of integrated resources, deployed more or less as a whole depending on how much interpretive support the local context provides.

Alternatively, if shallow parsing under GEP is specifically a matter of how much or which structure a listener predicts and builds, rather than a wholesale discounting of every operation that touches that structure once it exists, then the pattern of modulation should be graded rather than uniform, with semantic predictability affecting some components of syntactic processing more than others. Which components prove most sensitive is itself informative: a selective effect would indicate that semantic context does not simply reduce syntactic effort as a whole, but acts on particular computational demands, whether those tied to predicting and building structure, those tied to maintaining it in memory, or those tied to retrieving and integrating a dependent once its head is available. Adjudicating between a uniform and a graded pattern, however, requires estimating the influence of semantic context on several syntactic operations at once rather than on any one in isolation. In what follows, we describe an approach that models these operations concurrently and allows their sensitivity to semantic predictability to be compared directly.

## Materials and methods

### Participants and Procedure

We reanalyzed an open-source dataset of 61-channel EEG recordings obtained from 33 adult participants while they listened to an audio recording (∼12 minutes long) of Alice’s Adventures in Wonderland. Participants were between ages 18-28 and had normal hearing. Following the audiobook, participants completed a questionnaire about the story content to verify that they had paid attention and understood the story. For additional information regarding participant characteristics and recording parameters, we refer readers to the original manuscript from Brennan and Hale (B+H; Brennan & Hale, 2019). Four participants were excluded from further analysis for showing poor overall model fit (mean predictive correlation across the full predictor set at or below 0.005; see *Model estimation* below), leaving 29 participants for the primary analysis.

### Preprocessing

Raw data were preprocessed in the FieldTrip toolbox (Oostenveld et al., 2011) using the parameters described by B+H, consisting of high-pass filtering at 0.1 Hz, removal of ocular and other identified artifacts via Independent Component Analysis, and interpolation of high-impedance and noisy channels. Unlike B+H’s original sentence-epoched approach, data were retained as a single continuous recording per participant to support temporal response function estimation across the full audiobook rather than within individual sentences. Continuous data were resampled to 150 Hz following artifact correction and then band-pass filtered into the delta band (1-3 Hz) using a zero-phase FIR filter.

### Syntactic Predictors

We derived four measures of syntactic processing demand, each targeting a distinct computational operation. Dependency locality was quantified using the integration and storage cost metrics of Dependency Locality Theory (Gibson, 2000) as operationalized in recent fMRI work (Shain et al., 2022) and computed from dependency parses of the audiobook transcript using the Berkeley Natural Parser (Kitaev & Klein, 2018; Kitaev et al., 2019). Integration cost indexes the retrieval demand incurred when a word attaches to an earlier-occurring head, scaled by the number of new discourse referents (defined here as nouns and verbs) intervening between the dependent and its head; storage cost indexes the working-memory demand of maintaining currently unresolved syntactic dependencies, computed at each word as he number of syntactic dependencies that had been initiated by a preceding word but not yet completed; that is, the number of predicted heads or dependents still awaiting their partner at that point in the sentence, indexing the number of incomplete dependencies held in working memory as the sentence unfolds.

Syntactic surprisal was derived from a probabilistic context-free grammar rather than from a lexical language model, so that the resulting measure indexed prediction over hierarchical structure specifically rather than the broader mixture of statistical regularities that word-sequence surprisal conflates with structure. Constituency parses for each sentence in the transcript were also obtained using the Berkeley Neural Parser. Production-rule probabilities were estimated by maximum likelihood over the full set of rules observed across the audiobook, with additive smoothing applied to guard against zero-probability rules. Word-level syntactic surprisal was then computed via top-down attribution: for every internal node in a parse, its rule surprisal (the negative log probability of the rule that generated it) was assigned to the leftmost word dominated by that node, and a word’s total surprisal was the sum of all rule surprisals attributed to it in this way. This yields a measure of how unexpected the syntactic material introduced at a given word is, given the grammar’s statistics, without reference to lexical identity or sequential word statistics.

Closing-node count indexed the local structural work of completing a constituent, computed as the number of constituents in the parse tree for which a given word was the rightmost daughter. We selected closing-node count and structurally-derived surprisal as complementary rather than competing indices of syntactic processing, in part because prior work using node-count metrics has reached conflicting conclusions about which parsing algorithm, bottom-up, top-down, or left-corner, best predicts delta-band activity (Brennan et al., 2016; Coopmans et al., 2025; Rafferty et al., 2026; see also, Nelson et al., 2017 and Giglio et al., 2024 for compatible ECoG and fMRI findings, respectively), with outcomes plausibly depending on paradigm, modality, and analysis choices. Rather than adjudicate between competing parsing algorithms, we treated structural completion (closing-node count) and hierarchical prediction (surprisal) as two distinct computational demands with independent theoretical motivation, following the broader decomposition of syntactic processing described above. A schematic of syntactic variables can be seen in Figure 1. Full details of the grammar induction, smoothing, and attribution procedure, including the treatment of unary productions and multi-token constituents, are provided in the Supplementary Material.

**Figure 1:**
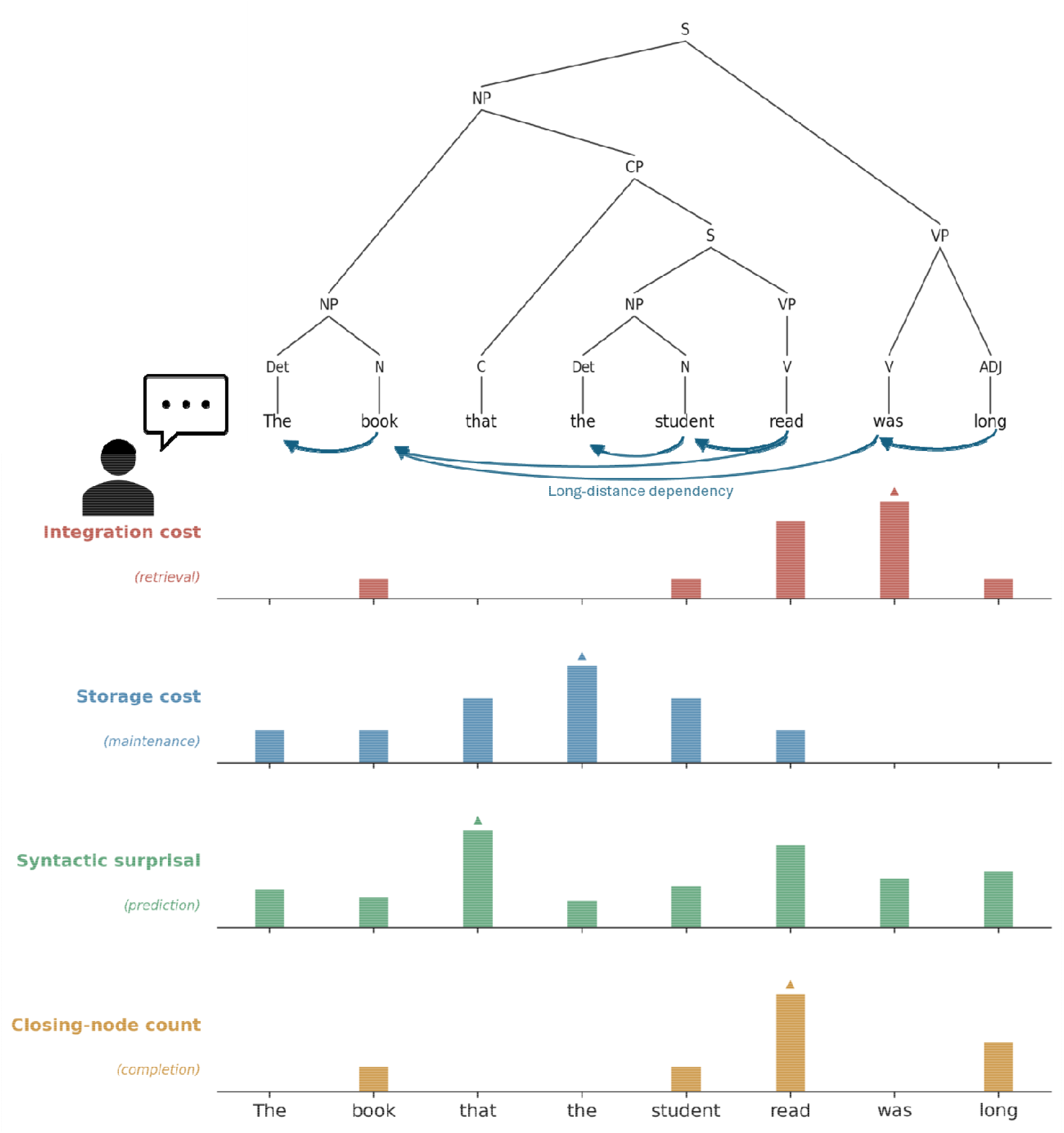
Schematic illustration of the four syntactic processing demands modeled in this study, shown for an example sentence containing a subject-modifying relative clause. The constituency tree (top—above the sentence) shows the hierarchical structure assigned to the sentence. Curved arrows (middle—below the sentence) marks the implemented backwards-looking dependency parse of the sentence. Each panel below plots the value of one syntactic predictor at each word, illustrating that the four measures index distinct computational operations and peak at different points in the sentence.

### Semantic Dissimilarity

Local lexical-semantic context was quantified as contextual dissimilarity, following the approach introduced by Broderick and colleagues (2018; 2019; 2022). Rather than static word vectors, we computed dissimilarity from contextualized word embeddings, so that a word’s representation reflected its specific usage rather than an average over all its possible senses. Embeddings were obtained from an intermediate layer of RoBERTa (Liu et al., 2019) for every content word in the transcript. For each word, dissimilarity was calculated as one minus the cosine similarity between that word’s embedding and the running mean embedding of all preceding words in the same sentence, such that dissimilarity increases as a word’s local semantic content diverges from its sentential context. Values were z-scored prior to predictor construction.

Broderick and colleagues showed that this measure elicits a centro-parietal negativity between roughly 200 and 600 ms that is diminished for time-reversed, unattended, and unintelligible speech, indicating that it indexes the semantic processing of comprehended speech rather than a purely distributional property of the stimulus. In subsequent work, semantic dissimilarity and lexical surprisal were shown to be only weakly correlated and to capture dissociable contributions to the neural response, consistent with their indexing predictive processing at semantic and lexical levels respectively. These properties make semantic dissimilarity well suited to the present design: it provides a validated, comprehension-sensitive index of how predictable a word is given its semantic context, and one that is separable from the structural predictors it is here crossed with, allowing its modulation of syntactic processing to be estimated without confounding semantic and syntactic sources of predictability.

### Predictor Construction

Continuous predictors were constructed at the EEG sampling rate (150 Hz) by placing an impulse at each word’s onset, with impulse height equal to that word’s value on the relevant measure. In addition to the four syntactic predictors and the semantic dissimilarity predictor described above, we included word onset (unit impulses marking every word) and word frequency (log-transformed corpus frequency) as control predictors. Predictors, along with their predictor category and operationalizations in the current work, are depicted in Table 1.

**Table 1:** Predictors entered into the temporal response function model. All eleven predictors were fit jointly in a single boosting model; the unique contribution of each was assessed by comparing this full model to a reduced model omitting that predictor (leave-one-out Δr). Two lexical control predictors account for onset-locked and lexical-access responses. Four syntactic predictors index distinct components of syntactic processing: retrieval and maintenance demands (integration and storage cost, from Dependency Locality Theory), prediction over hierarchical structure (syntactic surprisal, from a probabilistic context-free grammar), and local structural completion (closing-node count). One semantic predictor quantifies a word’s fit to its preceding context. Four interaction predictors, each the product of a z-scored syntactic predictor and z-scored semantic dissimilarity, test whether the neural response to a given syntactic operation depends on local semantic dissimilarity. All predictors were constructed as impulse trains time-locked to word onsets.

| Predictor | Category | Operational definition |
| --- | --- | --- |
| Word onset | Control | Unit impulse at each word onset; controls for the onset-locked evoked response. |
| Word frequency | Control | Log-transformed corpus frequency at each word onset; controls for lexical access demand. |
| Integration cost | Syntactic (retrieval) | Dependency Locality Theory integration cost: retrieval demand when a word attaches to an earlier head, scaled by intervening discourse referents. |
| Storage cost | Syntactic (maintenance) | Dependency Locality Theory storage cost: number of predicted-but-unrealized syntactic heads held in memory at each word. |
| Syntactic surprisal | Syntactic (prediction) | PCFG rule surprisal attributed top-down to the leftmost word of each constituent; indexes prediction over hierarchical structure. |
| Closing-node count | Syntactic (completion) | Number of constituents completed at a word (word is rightmost daughter); indexes local structural completion. |
| Semantic dissimilarity | Semantic | 1 - cosine similarity between a word's contextual embedding and the running mean of prior words in the sentence; increases with lower semantic fit to context. |
| Integration cost x Semantic dissimilarity | Interaction | Product of z-scored integration cost and z-scored semantic dissimilarity. |
| Storage cost x Semantic dissimilarity | Interaction | Product of z-scored storage cost and z-scored semantic dissimilarity. |
| Syntactic surprisal x Semantic dissimilarity | Interaction | Product of z-scored syntactic surprisal and z-scored semantic dissimilarity. |
| Closing-node count x Semantic dissimilarity | Interaction | Product of z-scored closing-node count and z-scored semantic dissimilarity. |

To directly test whether semantic predictability modulates each syntactic construct, we constructed four interaction predictors, one per syntactic measure, as the product of that measure’s z-scored impulse train and the z-scored semantic dissimilarity impulse train. Unlike prior work testing an analogous interaction by dichotomizing a continuous syntactic predictor into discrete high- and low-predictability categories (Slaats et al., 2024), our interaction predictors were constructed as continuous products of two z-scored measures, entered directly into the joint boosting model. This avoids the need to arbitrarily categorize a graded variable and estimates the interaction using the full range of variability in both measures simultaneously. This yielded eleven predictors in total: two lexical controls, four syntactic main-effect predictors, one semantic main-effect predictor, and four syntax-by-semantics interaction predictors.

Because interaction terms can be collinear with their constituent predictors, we assessed multicollinearity across all predictors prior to model fitting. Correlations among predictors were low (maximum absolute r = 0.41, between integration cost and its interaction with semantic dissimilarity), and all variance inflation factors fell well below the conventional threshold of 5 (maximum VIF = 1.28), indicating that the joint model’s coefficient estimates were not compromised by collinearity (Figure 2).

**Figure 2:**
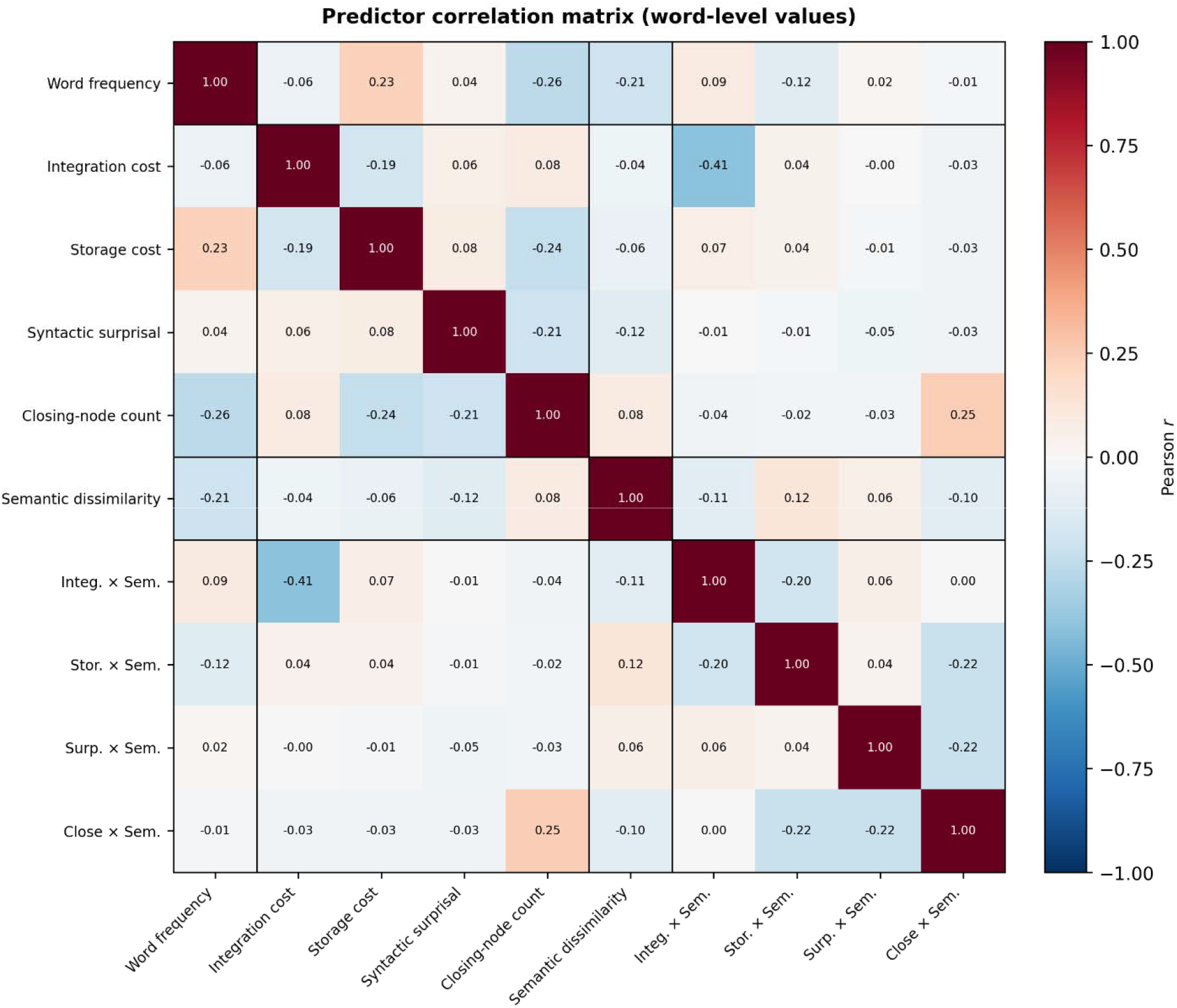
Correlation structure and multicollinearity among model predictors. Pearson correlations were computed across word-level predictor values; interaction terms were formed as the product of each z-scored syntactic predictor and z-scored semantic dissimilarity, exactly as entered into the model. Black lines separate predictor categories (lexical control, syntactic, semantic, and interaction terms). Correlations among predictors were generally low (maximum absolute off-diagonal r = 0.41, between integration cost and its interaction term). All variance inflation factors were well below the conventional threshold of 5 (maximum VIF = 1.28), indicating that multicollinearity, including between interaction terms and their constituent predictors, did not threaten the stability of the model coefficients. The word-onset predictor is omitted, as it is a constant unit impulse with no variance.

It is important to note that we did not include acoustic predictors such as the speech envelope or an acoustic onset spectrogram, which are commonly used in continuous TRF studies to control for low-level acoustic tracking. Two considerations motivated this choice. First, all of our linguistic predictors were sparse impulse trains time-locked to word onsets rather than continuous acoustic features, and the word-onset predictor included in every model absorbs the bulk of the onset-driven acoustic response that co-occurs with each word (Brodbeck et al., 2018). Second, and more importantly, our central hypotheses concern interaction terms, namely whether the neural response to a syntactic predictor depends on semantic dissimilarity. Because acoustic properties of the speech signal have no systematic relationship to the product of two z-scored linguistic measures, residual acoustic variance cannot plausibly account for a semantic-by-syntactic interaction, even if it were to contribute to a main effect.

### Model Estimation

Temporal response functions were estimated for each participant using the boosting implemented in Eelbrain (Brodbeck et al., 2023), with all eleven predictors entered into a single joint model. Boosting was performed over a lag window of -100 to 700 ms relative to word onset, with a 50 ms Hamming basis and ten-fold cross-validation. Model fit was quantified as the Pearson correlation between the model’s predicted response and the observed delta-band EEG signal on held-out data, averaged across folds, computed separately for each sensor. To isolate the unique contribution of each predictor, we additionally fit eleven reduced models, each omitting exactly one predictor from the full set. For each predictor, we computed Δr as the difference between the full model’s predictive correlation and the corresponding reduced model’s predictive correlation at each sensor; a positive Δr indicates that a predictor explains variance not accounted for by the remaining ten.

### Statistical Analysis

Group-level effects for each predictor’s Δr were evaluated using one-sample cluster-based permutation tests (10,000 permutations) across the scalp, testing whether Δr was reliably greater than zero (Maris & Oostenveld, 2007). This procedure was applied separately to each of the eleven predictors. To formally compare the relative magnitude of semantic modulation across the four syntactic constructs, we additionally computed each participant’s whole-scalp mean Δr for each of the four interaction predictors and submitted these values to a repeated-measures ANOVA with predictor identity as a within-subject factor. Significant omnibus effects were followed by pairwise paired-samples t-tests between all six predictor pairs, with Holm-Bonferroni correction for multiple comparisons. Because the cluster-based tests above use sensor selection informed by significance within the same data, this whole-scalp comparison was computed independently of any cluster-derived sensor subset, avoiding circularity in the magnitude comparison.

To characterize the direction and approximate time course of each significant interaction, we decomposed the temporal response function of each syntactic predictor into high- and low-semantic-dissimilarity components by adding and subtracting the corresponding interaction predictor’s response function, respectively. These conditional response functions were baseline-corrected over the pre-onset window and averaged across the sensors comprising each predictor’s significant interaction cluster. We treat these waveforms as a descriptive visualization of the direction and timing of the interaction effects established by the cluster-based analyses above, rather than as an independent inferential test.

## Results

### Model Fit and Participant Exclusion

We first assessed overall model fit for the full eleven-predictor boosting model, computed as the Pearson correlation between predicted and observed delta-band EEG activity, averaged across ten cross-validation folds and across sensors, for each participant. Whole-scalp mean predictive correlation across the sample of 33 participants ranged from -0.006 to 0.031 (median = 0.019), with peak per-sensor correlations ranging from 0.001 to 0.046 (median = 0.033). Four participants (S22, S25, S26, S44) showed whole-scalp mean predictive correlation at or below 0.005, indicating the model failed to capture reliable structure in their delta-band response. Because a whole-model fit this low reflects an absence of reliable stimulus-tracking in the recording rather than a selective failure of any one predictor and cannot be attributed to the linguistic variables of interest specifically, these participants were excluded from all subsequent analyses, leaving a final sample of 29. Results are depicted in Figure 3A and 3B.

**Figure 3:**
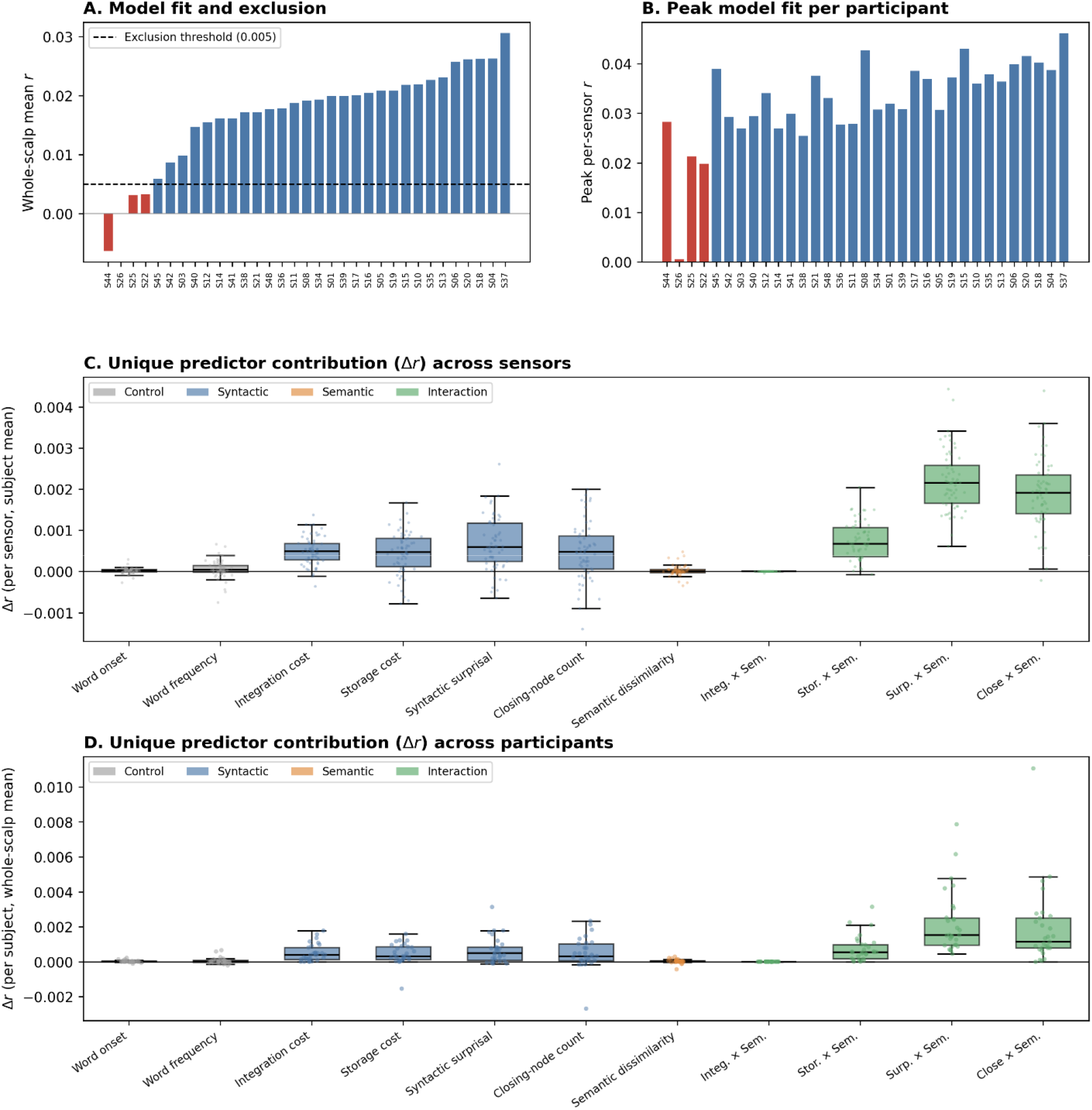
Model fit, participant exclusion, and unique predictor contributions. **(A)** Whole-scalp mean predictive correlation (r between predicted and observed delta-band EEG, averaged across ten cross-validation folds and all sensors) for each of the 33 participants, sorted in ascending order. The dashed line marks the exclusion threshold (r = 0.005); four participants falling at or below it (red) were excluded from all subsequent analyses. **(B)** Peak per-sensor predictive correlation for each participant, in the same order and coloring as (A), showing that retained participants’ best-fit sensors captured reliable signal. **(C)** Distribution of leave-one-out Δr across the 61 sensors for each predictor, averaged across the 29 retained participants; each point is one sensor, and boxes show the interquartile range and median. **(D)** Distribution of whole-scalp mean Δr across the 29 retained participants for each predictor; each point is one participant. This panel corresponds to the values entered into the repeated-measures comparison of interaction magnitude. In both (C) and (D), positive Δr indicates that a predictor explained variance not accounted for by the remaining ten predictors; predictors are colored by category (grey, lexical controls; blue, syntactic; orange, semantic; green, syntactic-by-semantic interactions).

### Syntactic and Semantic Main Effects

To evaluate whether each syntactic and semantic predictor explained variance in the delta-band signal beyond that captured by the remaining predictors, we computed, for each predictor, the difference in predictive correlation (Δr) between the full eleven-predictor model and a reduced model omitting that predictor, at every sensor. Group-level reliability of each predictor’s Δr was assessed using one-tailed, one-sample cluster-based permutation tests (10,000 permutations, cluster-forming threshold p < .05), separately for each predictor (Figure 3C and 3D).

All four syntactic predictors showed at least one significant positive cluster, indicating that each captured variance in the delta-band response not explained by the other ten predictors. Cluster-level significance was assessed by the summed t-statistic within each cluster (Maris & Oostenveld, 2007), reported below for the largest cluster of each predictor. Syntactic surprisal (cluster-*t* = 24.90, p < .001), integration cost (cluster-*t* = 15.87, p < .001), storage cost (cluster-*t* = 10.50, p = .002), and closing-node count (cluster-*t* = 6.56, p = .026) each yielded a reliable cluster. Because we did not conduct pairwise comparisons among these four main effects, we do not interpret differences in the magnitude or spatial extent of their clusters. Neither lexical control predictor showed a reliable cluster (word onset, cluster-*t* = 3.95, p = .080; word frequency, cluster-*t* = 1.74, p = .402), and semantic dissimilarity showed no significant cluster as a main effect, indicating that semantic predictability did not independently improve model fit once the four syntactic predictors and lexical controls were accounted for. Cluster info and TRF waveforms are depicted in Figure 4.

**Figure 4:**
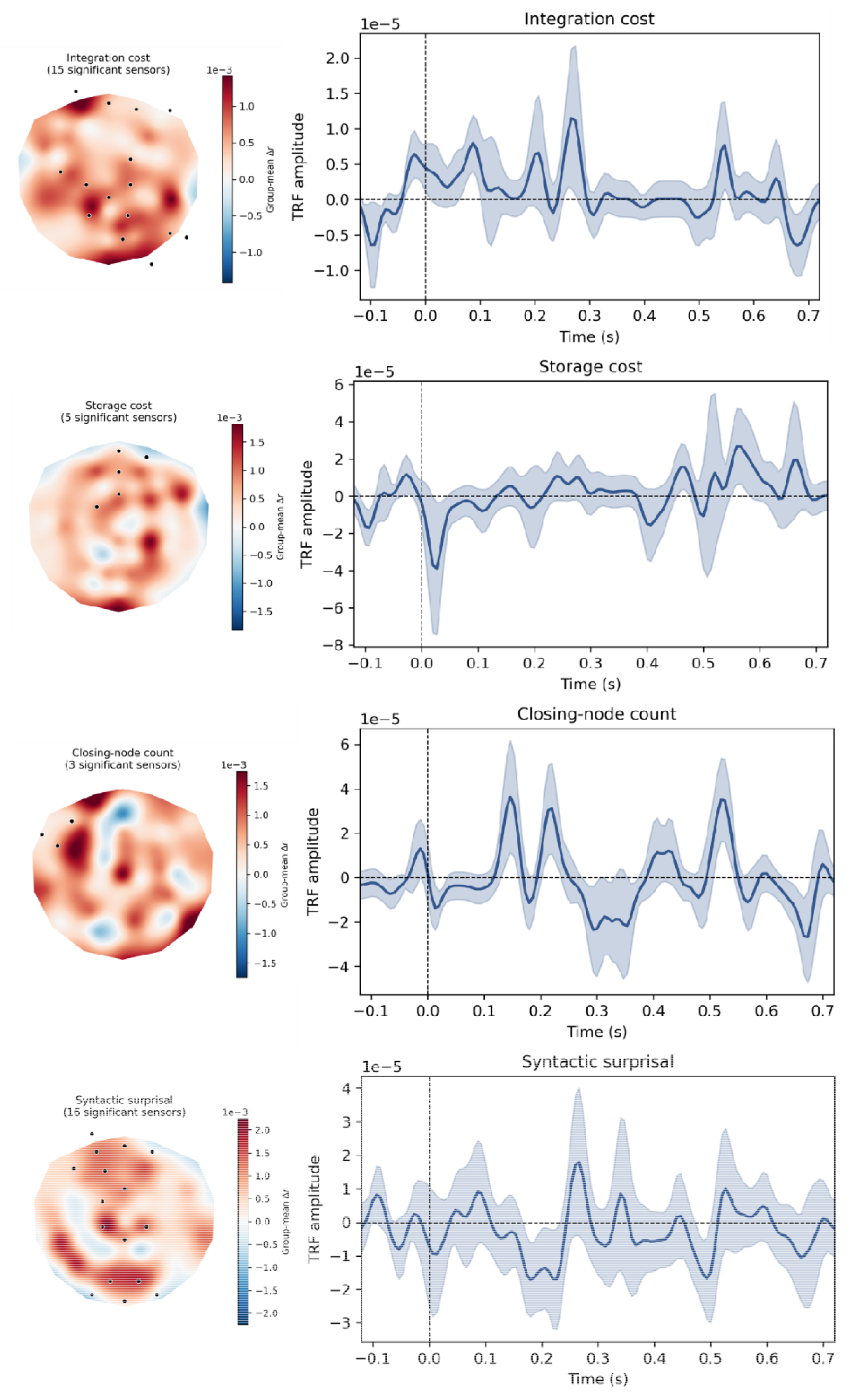
Syntactic main effects: significant electrodes and temporal response functions. For each of the four syntactic predictors, the left panel shows the group-mean Δr topography across the scalp, with electrodes belonging to significant cluster(s) marked in black (leave-one-out cluster-based permutation test, one-tailed, 10,000 permutations); the number of significant electrodes is given in each panel title. The right panel shows the corresponding group-mean temporal response function, averaged over that predictor’s significant electrodes and baseline-corrected over the pre-onset window (−100 to 0 ms); the shaded band denotes ±1 standard error of the mean across participants (n = 29). Positive Δr indicates that the predictor explained variance in the delta-band EEG signal not accounted for by the remaining ten predictors. All four syntactic predictors, integration cost, storage cost, syntactic surprisal, and closing-node count, showed significant positive clusters, confirming that each contributed uniquely to the model.

### Semantic Modulation of Syntactic Processing

The critical test of whether semantic predictability modulates the neural cost of syntactic processing came from four interaction predictors, each constructed as the product of a syntactic predictor and semantic dissimilarity, and each tested using the same cluster-based permutation procedure described above (Figure 5). Three of the four interactions showed significant positive clusters, indicating that the contribution of the corresponding syntactic predictor to the delta-band response depended reliably on local semantic dissimilarity. The interaction with syntactic surprisal was significant across the entire sensor array (61 of 61 sensors; cluster-*t* = 174.00, p < .001). The interaction with closing-node count was significant over a large majority of sensors (48 of 61; cluster-*t* = 127.77, p < .001), as was the interaction with storage cost (37 of 61; cluster-*t* = 78.15, p < .001). The interaction with integration cost showed no significant cluster, indicating no reliable evidence that semantic dissimilarity modulated this predictor’s contribution to the delta-band response. Importantly, this null result is not attributable to collinearity: although integration cost showed the highest correlation with its own interaction term of any predictor pair (r = −0.41), its variance inflation factor remained low (VIF = 1.28), and residual collinearity of this magnitude would if anything inflate rather than suppress an apparent effect.

**Figure 5:**
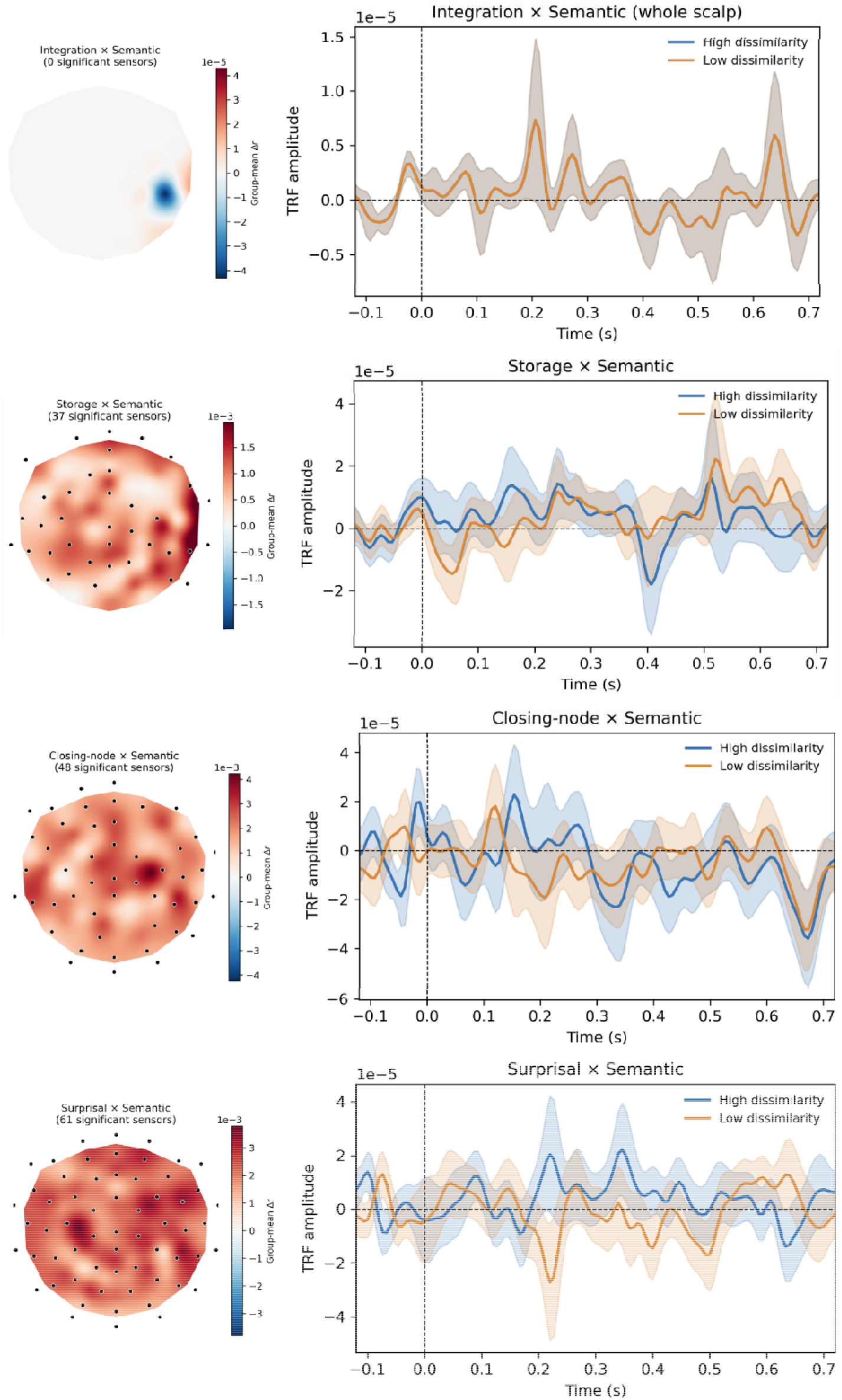
Semantic modulation of syntactic processing: significant electrodes and conditional temporal response functions. For each of the four syntactic predictors, the left panel shows the group-mean Δr topography for its interaction with semantic dissimilarity, with electrodes belonging to significant cluster(s) marked in black (leave-one-out cluster-based permutation test, one-tailed, 10,000 permutations); the number of significant electrodes is given in each panel title. The right panel shows the corresponding temporal response function decomposed by semantic dissimilarity: the response under high semantic dissimilarity (less predictable context; blue) and low semantic dissimilarity (more predictable context; orange), obtained by adding and subtracting the interaction response function from the syntactic predictor’s response function, respectively. Waveforms are averaged over the interaction’s significant electrodes and baseline-corrected over the pre-onset window (−100 to 0 ms); shaded bands denote ±1 standard error of the mean across participants (n = 29). The interactions with syntactic surprisal, closing-node count, and storage cost each showed significant positive clusters, indicating that the neural response to these predictors depended on local semantic dissimilarity; the interaction with integration cost showed no significant cluster, and its response functions are averaged over the whole scalp for illustration. These conditional waveforms are presented to characterize the direction and approximate timing of the interaction effects, not as an independent statistical test.

To characterize the direction and time course of these effects, we decomposed each syntactic predictor’s estimated temporal response function into high- and low-dissimilarity components, using the corresponding interaction term’s coefficients (Figure 5). For syntactic surprisal, the high- and low-dissimilarity response functions diverged most clearly between approximately 150 and 250 ms post-word-onset, with the low-dissimilarity condition showing a pronounced positive deflection largely absent in the high-dissimilarity condition. Closing-node count showed a similar, though temporally broader, divergence between dissimilarity levels. Storage cost showed a smaller and more temporally circumscribed divergence, consistent with its more limited spatial cluster. Integration cost showed no discernible separation between dissimilarity levels at any latency, consistent with the absence of a reliable interaction cluster for this predictor.

### Relative Magnitude of Semantic Modulation Across Syntactic Constructs

To formally evaluate whether the four syntactic constructs differed in the magnitude of semantic modulation they showed, rather than relying on cluster extent alone, we computed each participant’s whole-scalp mean Δr for each of the four interaction predictors and submitted these values to a repeated-measures ANOVA with predictor identity (integration cost, storage cost, syntactic surprisal, closing-node count) as a four-level within-subject factor. This analysis used whole-scalp averages rather than cluster-restricted sensors to avoid circularity, since cluster membership was itself determined from the same data.

There was a significant main effect of predictor identity, F(3, 84) = 14.02, p < .0001, indicating that the magnitude of semantic modulation differed reliably across the four syntactic constructs. Pairwise comparisons (paired-samples t-tests, Holm-Bonferroni corrected for six comparisons) resolved this effect into a graded hierarchy. Modulation of syntactic surprisal (M = .0022, SD = .0018) did not differ reliably from modulation of closing-node count (M = .0019, SD = .0022), t(28) = 0.53, p = .597, indicating these two predictors showed comparably strong semantic modulation. Both syntactic surprisal and closing-node count showed significantly greater modulation than storage cost (M = .00075, SD = .00074): surprisal versus storage cost, t(28) = 3.70, p = .003; closing-node count versus storage cost, t(28) = 3.28, p = .006. Storage cost, in turn, showed significantly greater modulation than integration cost (M ≈ 0, SD = .000004), t(28) = 5.43, p < .001, as did syntactic surprisal, t(28) = 6.71, p < .001, and closing-node count, t(28) = 4.66, p < .001, relative to integration cost. One-sample t-tests confirmed that whole-scalp mean Δr was reliably greater than zero for syntactic surprisal, t(28) = 6.71, p < .001, closing-node count, t(28) = 4.66, p < .001, and storage cost, t(28) = 5.42, p < .001, but not for integration cost, t(28) = -1.00, p = .837, where the mean Δr did not differ reliably from zero. Results are depicted in Figure 6.

**Figure 6:**
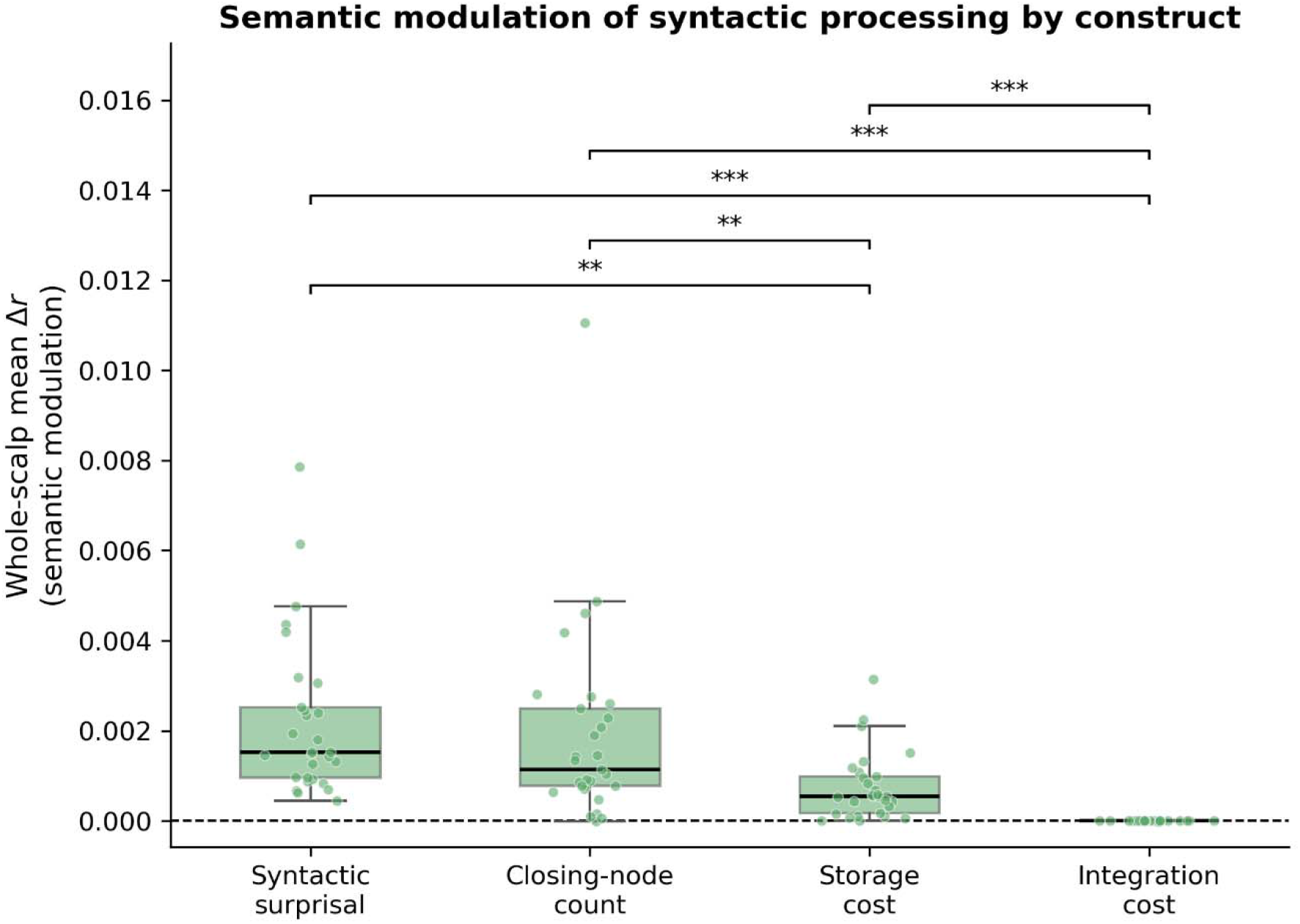
Magnitude of semantic modulation across the four syntactic constructs. Each point is one participant’s whole-scalp mean Δr for that construct’s interaction with semantic dissimilarity. Boxes show the median and interquartile range. The dashed line marks zero (no modulation). Brackets across the top of the figure denote Holm-Bonferroni-corrected pairwise comparisons (** p < .01, *** p < .001).

## Discussion

In this study, we used delta-band temporal response function modeling to evaluate whether local semantic predictability modulates the neural cost of syntactic processing during naturalistic speech comprehension, and whether this modulation is uniform across different syntactic operations or selective to particular ones. Our results support the latter. Semantic dissimilarity reliably modulated the neural response to syntactic surprisal and closing-node count, showed a smaller and more circumscribed modulation of storage cost, and showed no reliable modulation of integration cost at all. This graded pattern refines rather than simply confirms the good-enough processing (GEP) account: rather than a blanket down-weighting of syntactic effort whenever local semantics can support interpretation, our results suggest that semantic predictability selectively affects the anticipation and construction of syntactic structure, and the ongoing maintenance of that structure once built, while leaving the mechanics of retrieving and integrating an already-predicted dependent comparatively untouched.

This distinction matters for how “shallow parsing” under GEP should be understood mechanistically. A finding in which all four syntactic constructs were modulated equally would have been consistent with a genuinely undifferentiated efficiency strategy, in which the brain simply prioritizes syntax less across the board when semantics does the interpretive work. Our results instead suggest something more specific: when semantic context is highly predictable, listeners may generate less detailed structural predictions and maintain fewer competing structural commitments, but once a dependency is anticipated, integrating it proceeds in much the same way regardless of how predictable the broader context was. This is consistent with the possibility that GEP operates chiefly at the level of what structure gets anticipated and provisionally held in memory, rather than at the level of the retrieval operations that complete a dependency once it is available. It does not imply that listeners dispense with syntactic computation entirely in predictable contexts, nor that retrieval is somehow immune to processing demands more generally; rather, retrieval specifically may not be the locus at which contextual predictability exerts its influence.

Our findings converge with, and extend, two recent lines of work using similar multi-predictor TRF approaches. Weissbart and Martin (2024) found that syntactic and statistical predictors combine synergistically rather than additively in shaping neural dynamics, indicating that the two are not independently encoded and later summed but jointly determine the observed response. The interaction effects we report here are a direct instantiation of that synergy at the level of specific syntactic operations, showing where in the syntactic processing cascade that joint influence is concentrated. Slaats et al. (2024) reported that the delta-band response to syntactic closure is delayed by roughly 150-190 ms for high-relative to low-surprisal words, an interaction they estimated by dichotomizing surprisal into high and low categories and comparing the resulting node-count response functions. Our conditional response functions showed a comparable pattern for syntactic surprisal and closing-node count, with the clearest visual divergence between high- and low-dissimilarity conditions emerging in a similar window, around 150-250 ms post-word-onset. This convergence is notable for two reasons. First, our interaction was estimated continuously within a single joint model rather than via a median-split of the syntactic predictor as in previous work, suggesting the underlying effect is robust to how the interaction is operationalized. Second, the convergence was obtained using a different language, a different data set and corpus, and a structurally-derived rather than lexical surprisal measure, making it a useful cross-study replication of the general timescale at which semantic and statistical context shapes delta-band structure-building.

This timing pattern is broadly consistent with an oscillatory account in which the phase at which a linguistic unit arrives relative to an ongoing low-frequency oscillation carries information about that unit’s predictability, with predictable input arriving earlier, on a less excitable phase of the cycle (ten Oever & Martin, 2024; see also, Rafferty et al., submitted). Our design does not include phase information and speaks to this account only indirectly, but the present finding that semantic predictability shifts the estimated timing and magnitude of higher-level structural responses is compatible with the broader proposal that contextual predictability shapes not only lexical processing but the temporal dynamics of higher-level linguistic operations as well.

The selectivity of this modulation also bears on a distinction raised by Coopmans et al. (2022), who found that delta-band phrase tracking depends on the lexical-syntactic richness of an utterance’s input but not on whether the resulting structure composes into a coherent meaning. Our results are broadly compatible with, and add a further layer to, that picture. If phrase-level tracking indexes the building of structure from its input rather than the interpretability of its output, then the present findings suggest that semantic predictability acts on that same input-side process, shaping how much structure is anticipated and maintained, without correspondingly touching the comparatively mechanical operation of completing a dependency once the relevant input has already licensed it. Syntactic construction, in this view, remains sensitive to the local semantic context in which it unfolds, even though the eventual coherence of that context’s compositional output is not what delta-band tracking is reporting on.

These findings also sharpen how the two components of dependency locality relate to semantic context specifically. Integration and storage cost were developed within Dependency Locality Theory precisely to capture distinct demands, retrieval and maintenance respectively, and are typically reported together as complementary indices of working-memory load during sentence processing (Gibson, 2000; Shain et al., 2022). What has not previously been tested is whether these two demands are equally sensitive to the predictability of the local semantic context in which they occur. Our results indicate they are not: storage cost showed a modest but reliable interaction with semantic predictability, while integration cost showed none. This suggests that maintaining an unresolved dependency is more context-dependent than retrieving one once it becomes available, a distinction that is orthogonal to the retrieval/maintenance split DLT already draws and points to a further dimension along which the two components can be dissociated.

Taken together, these findings situate good-enough processing within the broader family of rational, noise-tolerant accounts of comprehension, of which it can be seen as one instance. Noisy-channel and rational-inference models hold that comprehenders perform Bayesian inference over a speaker’s intended message given an uncertain input and their prior expectations, weighting bottom-up structural evidence against top-down expectation rather than computing an exhaustive parse (Gibson et al., 2013; Warren & Dickey, 2021; Warren et al., 2025; Ryskin et al., 2025; Chen et al., 2026). On such accounts, a comprehender should invest less in predicting and building structure precisely when a strong semantic prior can carry the interpretation, since that is where structural computation is most redundant with information already available from context. This is broadly what our data show: semantic predictability most reduced engagement with the predictive and constructive components of syntactic processing, while leaving the retrieval of an already-committed dependency, an operation for which a semantic prior offers little pre-emptive benefit, comparatively unaffected. The connection is a natural one, since the phenomena that motivate good-enough processing—e.g., the acceptance of implausible or ungrammatical structures when a plausible interpretation is readily available—are among the same phenomena that noisy-channel models were developed to explain. Read this way, the graded dissociation we report does more than refine the good-enough account: it begins to specify where in the syntactic processing stream rational effort-allocation is concentrated, connecting a largely descriptive account of shallow parsing to a computational account of why, and where, it should arise.

### Limitations

Several limitations qualify these conclusions. First, our syntactic surprisal measure was derived via top-down attribution over a full-sentence constituency parse rather than from a strictly incremental parser, meaning it reflects the surprisal of the structure ultimately assigned to a sentence rather than a true prefix probability computed word by word; the correspondence between this measure and online prediction should be interpreted with this in mind. Second, the conditional response functions we report showed some structure prior to word onset, which we interpret as consistent with anticipatory processing, since our word-onset predictor and other lexical controls were included in the same model and should absorb variance attributable to acoustic onset per se. A stronger test of this interpretation would require paradigms in which the onset of predictive information can be more precisely controlled. Third, the sensor-averaged waveforms we report reflect a mixture of spatially distinct effects with likely different latencies, which may explain their oscillatory appearance. The topographic cluster analyses, rather than the sensor-averaged waveforms, should be treated as the primary evidence for the effects we report. Fourth, we did not explicitly test whether word duration, which is known to correlate with lexical surprisal and can itself be a cue to syntactic structure (Kuperman & Bresnan, 2012), contributed independently to the interaction effects we report; future work using duration-matched comparisons would help rule out this possibility. Finally, our findings are based on a single English-language audiobook and a comprehension sample of modest size. Replication across languages, genres, and larger samples will be necessary to establish the generality of the graded pattern we observe, particularly given evidence that even closely related languages can differ in the parsing strategies their grammatical structure invites.

### Future Directions

Other tenets of GEP beyond the modulation of syntactic processing cost, including the persistence of unresolved structural ambiguity in working memory and the conditions under which shallow parses lead to outright misinterpretation, remain outside the scope of the present study and would require different paradigms, such as garden-path or ambiguity-resolution designs, to address directly. Replicating the present graded pattern in other languages and genres, and testing whether the same maintenance/retrieval dissociation within dependency locality holds under other manipulations of context, would further establish how general this selectivity is. A natural further step would be to make the link to rational inference accounts explicit rather than interpretive. The present design establishes that semantic predictability selectively modulates syntactic processing, but it treats semantic dissimilarity as a single graded predictor rather than as a quantity within a formal inference model. A Bayesian extension could instead parameterize the comprehender’s prior directly, for instance by manipulating the strength or reliability of semantic context and asking whether the degree of structural down-weighting tracks the precision of that prior as a rational model would predict. Such a design would move from showing that structural computation is sensitive to semantic predictability toward testing whether that sensitivity is quantitatively rational and would allow the relative contributions of prior strength and bottom-up structural evidence to be estimated rather than inferred. Consistent with the broader literature on GEP, we do not intend the present findings as an endorsement of good-enough processing in its entirety. Rather, what they establish is that the neural cost of syntactic processing is not uniformly discounted under high semantic predictability but is instead selectively concentrated in the anticipation and construction of structure. This pattern sharpens rather than merely reiterates the claim that listeners prioritize efficiency during real-time language comprehension.

## Conclusion

Neural tracking of syntactic structure in the delta band is shaped by the local semantic context in which that structure is built, but not uniformly so. Using a temporal response function approach that modeled four distinct components of syntactic processing concurrently, we found that semantic predictability modulates the anticipation and construction of syntactic structure, exerts a smaller influence on its ongoing maintenance, and leaves the retrieval of an already-resolved dependency largely unaffected. These findings refine good-enough processing accounts of language comprehension by identifying which components of syntactic computation are, and are not, sensitive to the predictability of local meaning, and point toward multi-construct, interaction-based designs as a necessary tool for further specifying how the brain allocates syntactic effort during naturalistic language use.

