## Supplementary Materials for "Semantic predictability selectively modulates delta-band syntax tracking: evidence for good-enough parsing"

**Supplementary Material**

**S1. Detailed computation of syntactic surprisal and closing-node count**

This section provides the full procedure used to derive the two constituency-based syntactic predictors, syntactic surprisal and closing-node count, from the audiobook transcript. The main-text Methods summarize this procedure; here we give the level of detail required for reproduction.

**S1.1. Constituency parsing**

Each sentence in the transcript was parsed with the Berkeley Neural Parser (benepar; Kitaev & Klein, 2018), applied within a spaCy pipeline (en_core_web_md). Parsing produced a labeled constituency tree for every sentence. Where a transcript sentence was parsed into more than one sentence by the pipeline, each resulting sub-sentence was parsed and processed independently, and the resulting per-word values were concatenated in their original order.

**S1.2. Grammar induction and rule probabilities**

We estimated a probabilistic context-free grammar directly from the parsed audiobook rather than importing rule probabilities from an external treebank, so that the resulting surprisal values reflected the statistics of the stimulus itself. From every parse tree we extracted each internal production rule, defined as a left-hand-side nonterminal (a phrasal node label, e.g. NP) together with the ordered set of its immediate children's labels (e.g. NP → DT NN). Crucially, preterminal rules, that is, rules rewriting a part-of-speech tag as a specific word (e.g. NN → "book"), were excluded from the grammar. As a consequence, the surprisal measure is computed entirely from phrasal rule expansions and never has access to the identity of individual words. This is what licenses the interpretation of syntactic surprisal as an index of structural rather than lexical predictability: two constituents built from the same sequence of phrasal rules receive the same surprisal regardless of which specific words fill them.

The conditional probability of each rule was estimated by maximum likelihood with add-alpha (Laplace) smoothing over the observed rule inventory. For a rule with left-hand side *L* and right-hand side *R*,

*P(R | L) = (count(L → R) + alpha) / (count(L) + alpha * V_L)*

where count(L → R) is the number of times that rule occurred across the corpus, count(L) is the total number of expansions of *L*, V_L is the number of distinct right-hand sides observed for *L*, and alpha = 0.1. For any expansion of a left-hand side that was not among its observed right-hand sides, the smoothed probability was taken as alpha / (count(L) + alpha * V_L); the negative log of this quantity served as the surprisal of an unseen rule. Left-hand-side categories never observed at all (which did not occur in practice) were assigned a floor probability of 10^-10^.

**S1.3. Top-down surprisal attribution**

Following the top-down attribution scheme common to constituency-based surprisal measures in the naturalistic-comprehension literature, each internal rule's surprisal, defined as the negative log of its smoothed conditional probability, was assigned in full to the leftmost terminal (word) dominated by that rule's left-hand-side node. A given word's total syntactic surprisal was then the sum of the surprisal values of all rules attributed to it in this way. Because higher nodes in the tree contribute their surprisal to the leftmost word of their span, words that initiate large or low-probability structural constituents accrue higher surprisal.

We note that this measure is not incremental prefix-probability surprisal in the strict sense of Hale (2001), which would require computing the conditional probability of each word given all grammatical continuations consistent with the preceding prefix (for instance, via an Earley or left-corner parser run incrementally). Our measure instead reflects the summed rule surprisal of the structure ultimately assigned to a sentence, attributed top-down.

**S1.4. Closing-node count**

Closing-node count was computed for each word from the constituent spans returned by the parser, independently of the rule-probability computation above. For each sentence, we enumerated all constituents except the sentence-spanning constituent itself. Each constituent was assigned to the token at which it closed, that is, the final token of its span. A word's closing-node count was the number of constituents whose right edge fell on that word. Words at which several constituents close simultaneously, such as the final word of a right-branching embedded clause, therefore receive higher closing-node counts.

**S1.5. Alignment of parser output to the per-word data**

Because the parser's tokenization did not always match the word segmentation of the per-word EEG data table (for example, contractions and possessives such as "Alice's" were split by the parser into separate tokens), parser-derived values were aligned to the data table by a token-matching procedure. Tokens were compared after lowercasing and removal of non-alphanumeric characters. When a single data- table word corresponded to several parser tokens, the surprisal values of those tokens were summed and assigned to that word, and their closing-node counts were likewise summed; when a single parser token spanned several data-table words, its value was assigned to the first of those words and zero to the remainder. Sentences that could not be fully aligned by this procedure (a small minority) had their successfully aligned word positions retained, with any remaining unaligned surprisal positions filled with the corpus mean surprisal; closing-node counts for unaligned positions were set to zero.

**S1.6. Software**

Parsing used benepar (v0.2.0, benepar_en3 model) within spaCy (v3.8.14, en_core_web_md v3.8.0). Rule extraction, probability estimation, surprisal attribution, and closing-node counting were implemented in Python using NLTK's (v3.10.0) tree representation.

**S2. Detailed computation of dependency locality measures**

Integration cost and storage cost were derived from dependency parses of the transcript, following the operationalization of Dependency Locality Theory (Gibson, 2000) used by Shain and colleagues (2020) in their analysis of naturalistic story comprehension, an operationalization that has since been applied in subsequent naturalistic-comprehension work (e.g. Shain et al. 2022; Rafferty et al. 2026). The transcript was parsed with spaCy (en_core_web_md), and both metrics were computed from the resulting dependency structure in a single left-to-right pass over the tokens.

**S2.1. Discourse referents**

Both metrics depend on the notion of a discourse referent. Following the operationalization of Shain et al. (2020), we identified discourse referents from spaCy part-of-speech tags, distinguishing event-denoting from entity-denoting referents: nouns and proper nouns (POS tags NOUN, PROPN) were assigned a weight of 1, and verbs (POS tag VERB) a weight of 2 when they carried past- or present-tense morphology and 1 otherwise; all other tokens contributed 0. The heavier weighting of finite verbs reflects the assumption that tensed verbs introduce eventualities that impose greater integration demand than the introduction of a single entity. This referent-weighting scheme provided the best fit to neural data in the naturalistic fMRI work of Shain et al. (2020) and has been adopted in subsequent studies of naturalistic comprehension.

**S2.2. Storage cost**

Storage cost was computed at each token as the number of syntactic dependencies that had been opened but not yet resolved at that point in the sentence. Concretely, as the parser's token sequence was traversed left to right, a dependency was counted as open when a token's head lay to its right (that is, the head had not yet been encountered). At each token, the set of open dependencies was restricted to those whose head still lay strictly ahead of the current position, and storage cost was the size of this set. Storage cost therefore increases across regions of a sentence in which multiple dependencies are simultaneously pending resolution, such as the interior of a center-embedded or pre-head-modified constituent, and falls as those dependencies are discharged.

**S2.3. Integration cost**

Integration cost was computed at each token that completed a dependency with an earlier head, that is, a token whose head lay to its left. For such a token, we summed the discourse-referent weights (Section S2.1) of the tokens occurring strictly between the head and the dependent. A dependency completed across a span containing several intervening referents therefore incurred a higher integration cost than one completed locally, capturing the core locality prediction of DLT that longer dependencies are more costly to integrate.

Two additional choices, also part of the operationalization we followed, governed this computation. First, modifier dependencies, comprising adjectival modifiers, adverbial modifiers, nominal adverbial modifiers, numeric modifiers, determiners, and possessives (spaCy dependency labels amod, advmod, npadvmod, nummod, det, poss), were assigned an integration cost of zero, on the assumption that the attachment of such local modifiers does not incur the retrieval demand associated with integrating arguments and heads. Second, coordinated structures received special treatment: rather than summing referent weights across all conjuncts of a coordination occurring within the integration span, we took the maximum conjunct weight, so that a coordination contributed to integration cost according to its heaviest single conjunct rather than the sum of its parts. This avoids over-penalizing the integration of coordinated material, whose conjuncts are structurally parallel rather than sequentially dependent.

**S2.4. Software**

Dependency parsing used spaCy (v3.8.15, en_core_web_md v3.8.0). Integration and storage cost were computed in Python from the resulting parse.

**S3. Detailed computation of semantic dissimilarity**

Semantic dissimilarity was computed per content word following the approach of Broderick et al. (2018), using contextualized word embeddings in place of the static vectors used in the original formulation.

**S3.1. Contextual word embeddings**

Each sentence was tokenized and passed through RoBERTa (roberta-base), and per-word embeddings were taken from the model's eighth hidden layer. Because the tokenizer segments words into byte-pair-encoding subword units, the subword embeddings belonging to each word were mean-pooled to yield a single embedding per word; special tokens were excluded from pooling. Embeddings were computed sentence by sentence, so that each word's representation reflected its context within its own sentence.

**S3.2. Running-context dissimilarity**

For each word i beyond the first in its sentence, semantic dissimilarity was computed as one minus the cosine similarity between that word's embedding and the mean embedding of all preceding words in the same sentence:

dissimilarity_i = 1 - cos( e_i , mean(e_1, ..., e_{i-1}) )

where e_i is the layer-8 embedding of word i and the running context mean is taken over every word preceding i within the sentence. Dissimilarity therefore increases as a word's contextual representation departs from the average representation of the context that preceded it. The first word of each sentence, having no prior within-sentence context, was assigned the sentence's mean dissimilarity. This formulation corrects the sign of the similarity term relative to an earlier, unpublished computation, such that higher values correspond to greater semantic divergence from context.

**S3.3. Software**

Contextual embeddings were obtained from RoBERTa (roberta-base) via the Hugging Face Transformers library (v5.8.1), with PyTorch (v2.12.0) as the backend.
